# A Multivariate Approach to Understanding the Genetic Overlap between Externalizing Phenotypes and Substance Use Disorders

**DOI:** 10.1101/2022.09.27.509777

**Authors:** Holly E. Poore, Alexander Hatoum, Travis T. Mallard, Sandra Sanchez-Roige, Irwin D. Waldman, Abraham A. Palmer, K. Paige Harden, Peter B. Barr, Danielle M. Dick

**Author notes:** These authors jointly supervised this work (Peter B. Barr, Danielle M. Dick). Correspondence should be addressed to Holly E. Poore, 671 Hoes Lane W, Piscataway, NJ 08854.

## Abstract

Substance use disorders (SUDs) are phenotypically and genetically correlated with each other and with other psychological traits characterized by behavioral undercontrol, termed externalizing phenotypes. In this study, we used Genomic Structural Equation Modeling to explore the shared genetic architecture among six externalizing phenotypes and four SUDs used in two previous multivariate GWAS of an externalizing and an addiction risk factor, respectively. Using a preregistered set of criteria, we first evaluated the performance of five confirmatory factor analytic models, including a common factor model, alternative parameterizations of two-factor structures, and a bifactor model. We used a combination of model fit, factor reliability, and model characteristics to adjudicate among the models. We next explored the genetic correlations between factors identified in these models and other relevant psychological traits. We found that a common factor model, in which all externalizing phenotypes and SUDs were influenced by a single dimension of genetic risk best characterized the relationships among our phenotypes. Although two two-factor models also performed well, we found that the factors in those models were very highly correlated with each other (*r*_gS_ > .87) and similarly genetically correlated with external criteria, suggesting they did not represent meaningfully distinct dimensions. Results from this study can be used to inform future efforts to characterize genetic liability for broad externalizing as well as specific externalizing phenotypes.

## Introduction

Substance use disorders (SUDs) are associated with substantial cost to affected individuals, their families, and society at large^1–3^. Twin and family studies estimate the heritability of individual SUDs, including alcohol^4^, cannabis^5^, and opioid use disorders^6^, to be around 50%, with a large portion of the heritability for each disorder shared across different substances^6^. SUDs co-occur with other forms of psychopathology, personality, and behavioral traits^7^, most notably with disorders and traits characterized by under-controlled or impulsive action, often termed *externalizing phenotypes*^8,9^. These phenotypes include psychiatric disorders, such as attention deficit hyperactivity disorder (ADHD), conduct disorder (CD), oppositional defiant disorder (ODD), and antisocial personality disorder (ASPD)^10–12^, as well as personality and behavioral traits like risk taking^13^, aggression^14,15^, lack of constraint^9^, and antagonism^16,17^. Mirroring phenotypic associations, SUDs are also genetically correlated with externalizing phenotypes^18–21^.

Evidence of strong genetic correlations among SUDs and externalizing phenotypes suggests that further investigation into their shared genetic architecture is warranted. Recent advances in statistical genetics methods allow us to leverage summary statistics from well-powered genome-wide association studies (GWAS) to model the degree to which genetic influences on individual, related phenotypes operate through broad latent factors that represent the variance shared among these phenotypes. Specifically, Genomic Structural Equation Modeling (Genomic SEM)^22^ provides a flexible framework to apply SEM techniques to genetic covariance matrices derived from Linkage Disequilibrium score regression^23^. This allows researchers to formally test theoretical models of the shared genetic architecture of genetically correlated traits.

Two recent large-scale studies used Genomic SEM to examine the genetic architecture of externalizing phenotypes (N = 1,373,240)^13^ and SUDs (N = 1,019,521)^24^, respectively. Both identified a single, underlying latent factor that captured the majority of the variance in the outcomes. A GWAS of liability to externalizing, operationalized as a latent factor indicated by attention deficit/hyperactivity disorder (ADHD), problematic alcohol use, cannabis initiation, smoking initiation, age at first sexual intercourse, number of sexual partners, and general risk tolerance, identified 579 conditionally independent loci that largely operated through the latent factor^13^. Similarly, a GWAS of liability to addiction, indexed by problematic alcohol use, problematic tobacco use, cannabis use disorder, and opioid use disorder, identified 19 independent loci^25^.

Results from these studies support the use of broad latent factors to interrogate the shared genetic etiology of externalizing phenotypes and SUDs. However, the question remains as to whether SUDs and externalizing are influenced by distinct, but related, dimensions of risk, or whether they reflect the same underlying continuum of risk. In other words, is there specific genetic risk shared across SUDs, that is not also shared with other externalizing outcomes? To address this question, we applied Genomic SEM to six externalizing phenotypes and four SUDs previously included in the factor models of externalizing^13^ and addiction liability^24,25^. We tested a series of preregistered, *a priori* specified models, guided by results from the phenotypic factor analysis literature. Previous studies have produced inconsistent results with respect to the placement of SUDs in the overall structure of psychopathology. Many studies include SUDs as part of a broad externalizing factor^26–29^, whereas others model it as a separate, but correlated, factor^11,30–32^. We thus tested five alternative confirmatory factor analytic models, including a common factor model and alternative parameterizations of a two-factor structure to capture the genetic covariance among externalizing phenotypes and SUDs (Figure 1a-1e). We followed up these analyses with exploratory factor analysis to evaluate alternative, data-driven clusters of phenotypes. Finally, we estimated genetic correlations between the factors identified in our models and a variety of other relevant traits to further characterize the extent to which these dimensions are differentially related to other outcomes.

**Figure 1:**
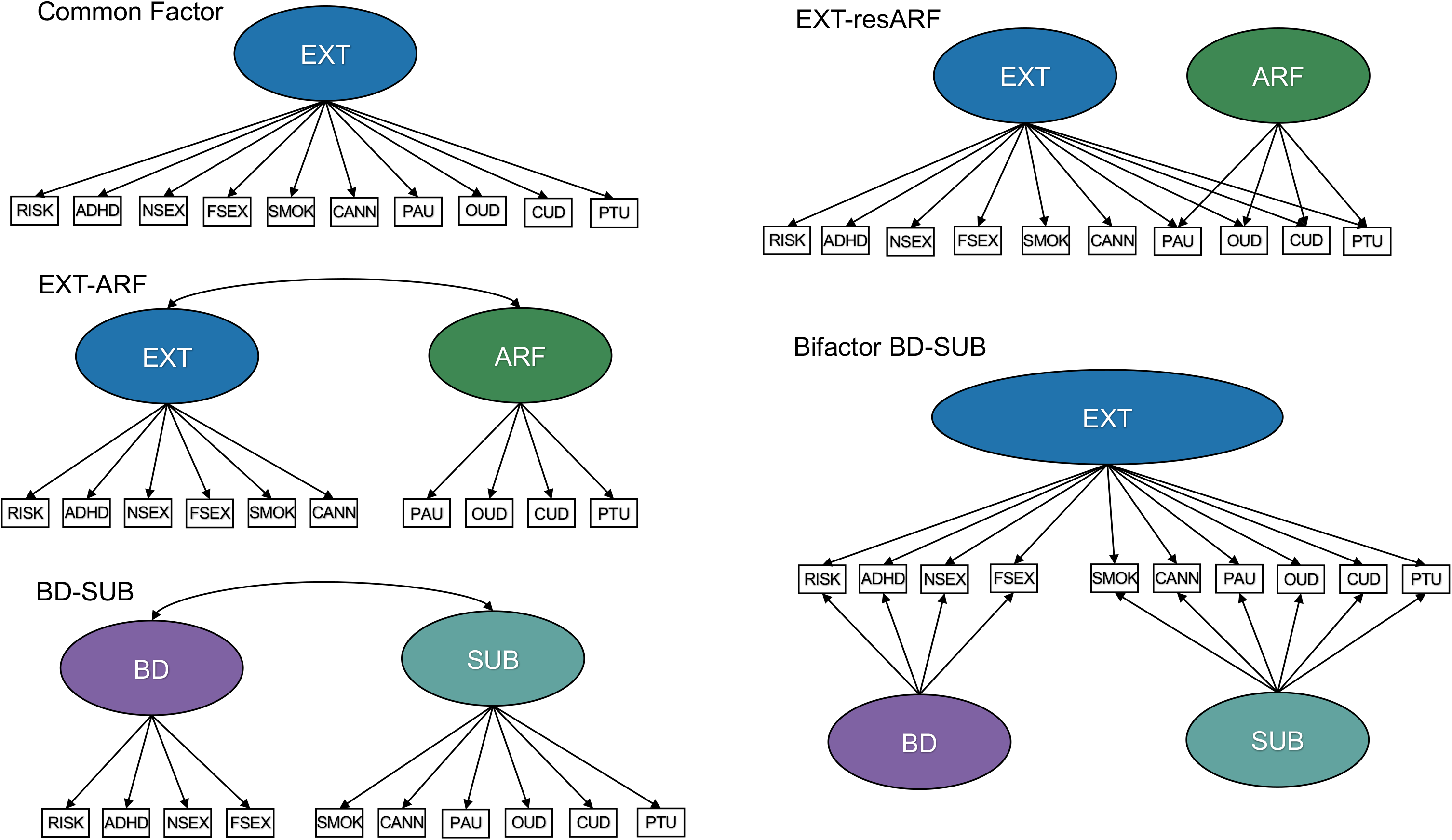
Confirmatory factor analytic models tested. Path diagrams of all confirmatory models tested (a) Common Factor, (b) EXT-ARF Model, (c) BD-SUB Model, (d) EXT-resARF Model, and (e) Bifactor BD-SUB Model.

## Methods

### Samples

Following our preregistered analysis plan (https://osf.io/v8q2y/), we used summary statistics from well-powered GWAS previously included in investigations of externalizing^13^ and the addiction risk factor^24^. We included the following 10 phenotypes: general risk tolerance^33^, number of sexual partners^33^, reverse coded age at first sexual intercourse^33^, attention deficit hyperactivity disorder^34^, smoking initiation^35^, cannabis initiation^36^, problematic tobacco use^24^, problematic alcohol use^37^, opioid use disorder^38^, and cannabis use disorder^21^. See Supplementary Table 1 for more information about each phenotype. As previous Genomic SEM studies of externalizing and addiction liability both included similar indicators for problematic alcohol use (N ~ 150K in Karlsson Linnér and colleagues^13^; N ~ 430K in Hatoum and colleagues^24^), we only retained the larger meta-analysis of problematic alcohol use. Following previous investigations using these GWAS summary statistics, we retained variants with minor allele frequency > .01 and INFO score > .70 for opioid use disorder and >.90 for all other phenotypes.

### Statistical Analyses

We used Genomic SEM^22^ to conduct all analyses. Genomic SEM is a recent statistical method that provides a flexible framework for applying SEM techniques to GWAS summary statistics, which allows for more accurate modeling of multivariate genetic covariance matrices.

In the present study, we first estimated the genetic correlations among the ten indicators in our proposed model. Next, we tested a series of *a priori* specified models to investigate the genetic architecture of externalizing phenotypes and SUDs. We tested the following models:

1. Common Factor Model: an externalizing factor onto which all externalizing phenotypes and SUDs load
2. EXT-ARF Model: a two-correlated factors model in which externalizing phenotypes load onto an externalizing factor (EXT) and SUDs load on an addiction risk factor (ARF)
3. BD-SUB Model: a two-correlated factors model in which all substance use phenotypes load onto a substance use factor (SUB)^39^ and remaining phenotypes comprise a behavioral disinhibition factor (BD)
4. EXT-resARF Model: a model in which all phenotypes load on an externalizing factor (EXT) and SUDs load onto a residual ARF
5. Bifactor Model: a model in which liability to externalizing is captured by a general factor onto which all phenotypes load and residual liability to behavioral disinhibition (BD) and substance use (SUB) is captured by two specific factors.

We assessed the performance of each alternative models using a combination of the following metrics:

1. Goodness-of-fit statistics, including the Comparative Fit Index (CFI), the Akaike Information Criterion (AIC) and the Standardized Root Mean Square Residual (SMSR), which provide absolute and relative indices of model fit. CFI and SRMR are absolute fit indices, with values greater than .95 and below .05, respectively, indicating excellent fit^40^.
2. Magnitude, median, and standard deviation of the indicators’ loadings on their respective factors in each alternative model^41^. If factors are representative of their indicators, indicators’ loadings will be substantial as indicated by their individual and average magnitude. If factors represent unidimensional constructs, they should be similarly defined by their constituent indicators^10,42^. This is particularly important as our first aim is to determine the degree of dimensionality in the previously identified externalizing phenotypes and SUDs. We thus compare the standard deviations of indicators’ loadings, which should be low in cases where factors are similarly represented by their indicators.
3. Structural validity will be assessed using construct replicability (H)^43^ and latent variable reliability (omega)^44^. H conceptually reflects the extent to which a latent variable is represented by its indicators (items). It ranges from 0-1 and increases as a function of the magnitude of factors loadings and number of indicators on each item. Although inherently arbitrary, values of greater than .70 indicate adequate replicability^43^. For latent variable reliability in the common factor model, we will use omega total (ω_t_), which reflects the percentage of total variance accounted for by the single latent construct. For the correlated factors models, we will use omega subscale (ω_s_), which reflects the percentage of variance the latent variable accounts for in its indicators. Values greater than .75 indicate adequate reliability^44^.
4. Model characteristics, such as standardized factor loadings that are out of bounds (i.e., > |1|), not in the predicted direction, not significantly different from zero, or have large standard errors, will be used to judge whether models are tenable^41^.
5. Sensitivity of the factors to their indicators, which will be assessed by dropping one indicator in the model at a time and judging the extent to which this changes the factor loadings and their standard errors of other indicators as well as the factor correlations in the EXT-ARF and BD-SUB 2CF Models.

To explore a wider range of potential factor solutions, we next conducted exploratory factory analyses (EFA) using the *stats* R package^45^. We tested 2-4 factor solutions using oblique (i.e., correlated) and orthogonal (i.e., uncorrelated) rotations. We tested solutions with a maximum of 4 factors to ensure factors would have more than 2 indicators. We used the genetic covariance matrix of odd chromosomes for the EFAs and then fit CFAs, replicating the exploratory results, using the genetic covariance matrix of even chromosomes.

Finally, we conducted genetic association analyses, estimating the zero-order genetic correlations between factors identified in the best-performing models and a wide range of preregistered phenotypes (https://osf.io/v8q2y/) from the domains of personality, risk taking, physical health, psychiatric traits and disorders, anthropometric traits, cognitive traits, socioeconomic status, and reproductive health. A full description of these phenotypes is provided in the Supplementary Information. These genetic association analyses served two purposes: 1) to explore the genetic relationships between our latent construct(s) and other relevant traits and 2) to compare patterns of genetic correlations in the best-performing two-factor models. This latter purpose allowed us to quantify the degree to which these factors provided meaningfully distinct information about genetic risk for externalizing phenotypes and SUDs. As a further test of this question, we fit models in which the correlations between the two factors and an individual external criterion variable were constrained to be equal and observed the resulting change in χ^2^. We note, however, that χ^2^ difference tests are very sensitive to small changes, especially in the presence of a large sample size, and interpret these results with caution. We used a Bonferroni corrected p-value < .05 to judge statistical significance.

## Results

### Model Fitting

We first estimated the genetic correlations among our ten proposed indicators. The correlations between SUDs and externalizing phenotypes varied substantially between SUDs (Figure 2). Of the four SUDs, cannabis use disorder was most strongly correlated with the externalizing phenotypes (*r*_gS_ ranged from .71 [age at first sex] to .72 [smoking initiation]) whereas problematic tobacco use was least strongly correlated with externalizing phenotypes (*r*_gS_ ranged from −.06 [cannabis initiation] to .45 [ADHD]).

**Figure 2:**
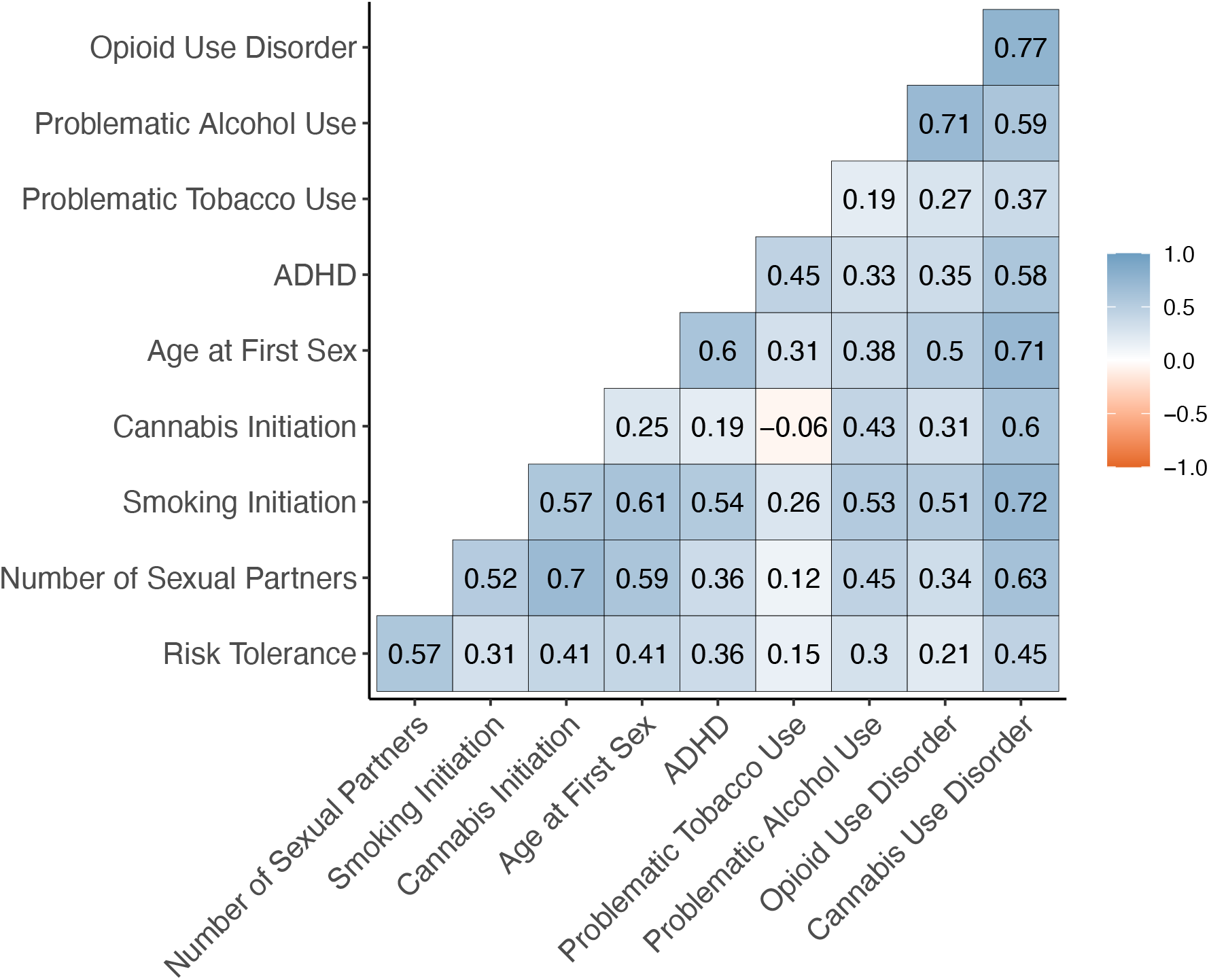
Genetic Correlations among Externalizing Phenotypes and SUDs

Given that SUDs were at least moderately correlated with most externalizing phenotypes, we proceeded with our prespecified set of confirmatory factor analytic models. See Table 1 for model fit and Table 2 for factor loadings and reliability indices. In addition to the confirmatory factor analyses presented below, we ran exploratory factor analytic (EFA) models with 2-4 factors. No clear pattern emerged from the EFAs, the results of which are reported in Supplementary Material (Supplementary Tables 2-3).

**Table 1.** CFA Model Fit Statistics

| **Model** | **$\chi^2$ (df)** | **AIC** | **CFI** | **SRMR** |
| --- | --- | --- | --- | --- |
| Common Factor Model (Model 1) | 1708 (35)<sup>***</sup> | 1748 | 0.86 | 0.10 |
| EXT-SUD (Model 2) | 1632 (34)<sup>***</sup> | 1674 | 0.87 | 0.10 |
| BD-SUB (Model 3) | 1390 (34)<sup>***</sup> | 1432 | 0.89 | 0.10 |
| EXT-residual SUD (Model 4) | 1684 (31)<sup>***</sup> | 1732 | 0.86 | 0.09 |
| BD-SUB Bifactor Model (Model 5) | 183 (24)<sup>***</sup> | 245 | 0.99 | 0.06 |

**Table 2.**
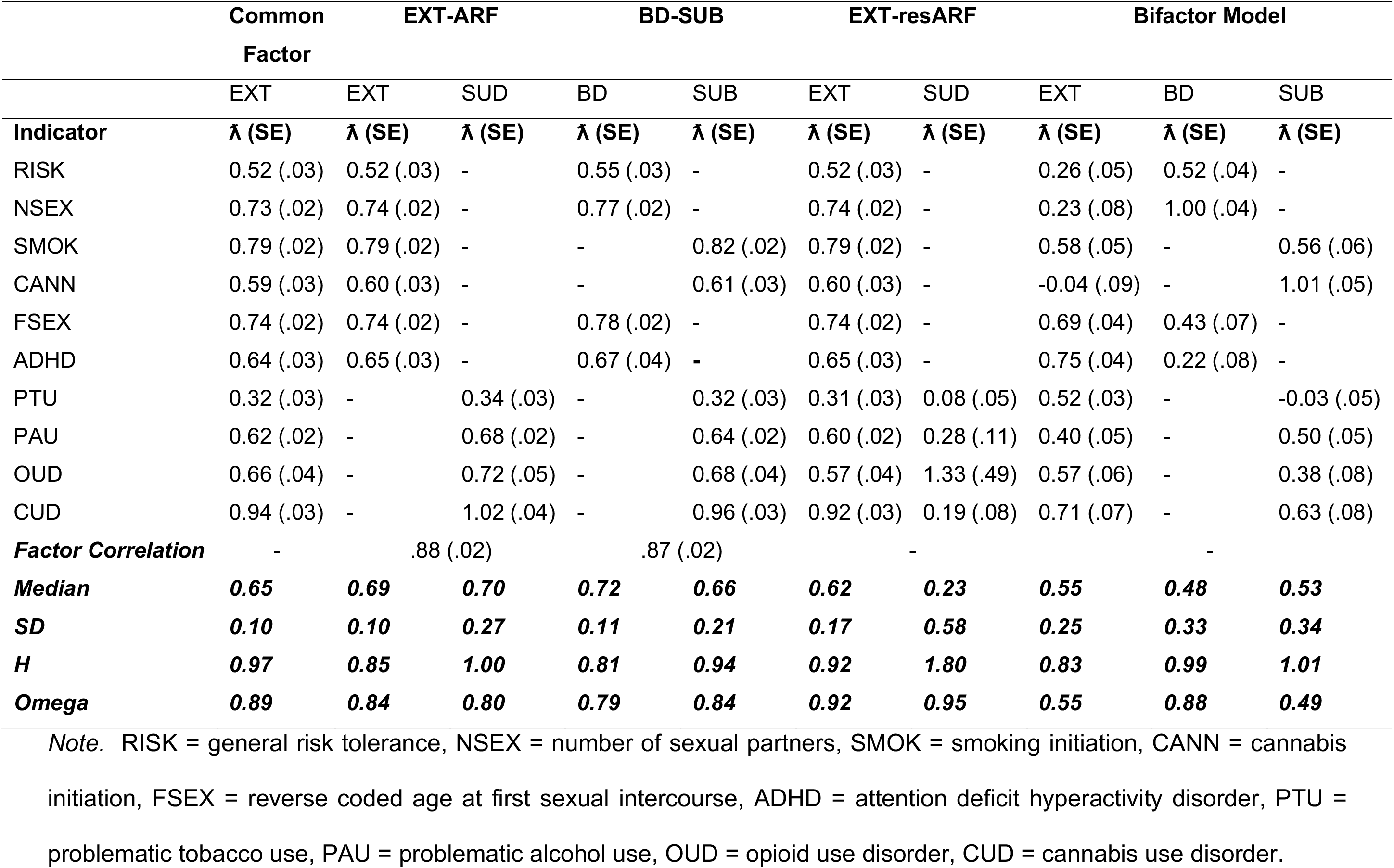
Standardized Factor Loadings & Factor Reliability Indices for CFA Models

#### Model 1: Common Factor Model

Fit statistics for the Common Factor model, in which all externalizing phenotypes and SUDs loaded onto a single factor, did not meet our prespecified threshold for good model fit. Nevertheless, the standardized factor loadings of the indicators were high (median ƛ = .65) and uniform (SD of ƛ = .10), indicating that the common factor was well and similarly defined by its indicators. In further support of this, H and ω_t_ were high (.97, and .89, respectively).

#### Model 2: Two-Correlated Factors Model with Externalizing and Addiction Risk Factors (EXT-ARF)

We next fit the EXT-ARF model, in which externalizing phenotypes loaded onto an externalizing (EXT) factor and SUDs loaded on an addiction risk factor (ARF). Fit statistics for this model also did not meet thresholds for good fit, although fit was somewhat improved relative to the Common Factor model. The median standardized factors loadings were high (ƛ = .66), although the standard deviation of the loadings on the ARF factor were somewhat higher (SD of ƛ = .27), indicating greater variability in the degree to which these factors were represented by their indicators. The EXT and ARF factors were very strongly genetically correlated (*r*_gS_ = .88), suggesting they reflect largely overlapping dimensions of genetic risk. H and ω_s_ were high for both factors, although it is important to note that the H for the ARF factor is uninterpretable given that cannabis use disorders’ loading exceeded 1.

#### Model 3: Two-Correlated Factors Model with Substance Use and Behavioral Disinhibition Factors (BD-SUB)

Fit statistics for the BD-SUB model, in which all substance use phenotypes loaded onto a substance use (SUB) dimension and remaining phenotypes loaded onto a behavioral disinhibition factor (BD) were improved relative to the Common Factor and EXT-ARF models, but still did not meet threshold for good fit. The median factor loadings were high (ƛ = .72), although the variability of loadings on the SUB factor was somewhat high (SD of ƛ = .21). The BD and SUB factors were strongly correlated (*r*_gS_ = .87), again suggesting that they represent similar dimensions of risk. H and ω_s_ were high for both factors.

#### Model 4: Externalizing Factor with residual Addiction Risk Factor (EXT-resARF)

The EXT-resARF model, in which all phenotypes loaded onto an externalizing (EXT) factor and SUDs loaded additionally onto a residual addiction risk factor (resARF), did not meet the threshold for good fit. The median loading on the residual ARF factor was low (median ƛ = .23) and the loadings were variable (SD of ƛ = .58). In addition, the residual variance of the OUD indicator was negative.

#### Model 5: Bifactor Model with Externalizing General Factor and specific Behavioral Disinhibition and Substance Use Factors (Bifactor BD-SUB)

Finally, we fit the BD-SUB Bifactor model, in which all phenotypes loaded on a general externalizing factor and residual variation among the phenotypes was captured by behavioral disinhibition and substance use specific factors. This model fit the data well based on fit statistics, but several indicators had negative residual variances or nonsignificant or negative loadings on the general and specific factors, likely reflecting overfitting. In addition, the median standardized factors loadings of the general and specific factors were low, and the standard deviations were high. H was high for most factors, although this is strongly influenced by the loadings greater than 1.0. ω was below the threshold of good reliability for the general factor and substance use specific factor (.55 and .49, respectively).

#### Summary of Model Fitting

Across all models, problematic tobacco use had the lowest loadings on its respective factors (ƛ ranged from .32 - .34 in the Common Factor, EXT-ARF, and BD-SUB models) and cannabis use disorder had the highest loadings (ƛ ranged from .94 – 1.02 in the Common Factor, EXT-ARF, and BD-SUB models). We tested a series of models dropping the problematic tobacco use indicators to test the impact this indicator had on model fit and found that model fit was not substantially impacted (see Supplementary Table 4).

No models met our preregistered criteria for good fit based on model fit indices. Nevertheless, we evaluated each model against the other evaluation criteria outlined in the methods (i.e., magnitude, median, and standard deviation of the indicators’ loadings, structural validity, and model characteristics) to decide which models to carry forward into subsequent analyses. We determined that the EXT-resARF and Bifactor BD-SUB models were untenable given evidence of low factor loadings, high standard errors, poor reliability, and negative residual variances. The three remaining models, the Common Factor, EXT-ARF and BD-SUB models, all had high loadings, good reliability, and no evidence of concerning model characteristics (i.e., high standard errors or negative residual variances). The factors in both the EXT-ARF and BD-SUB models were very highly genetically correlated, suggesting that the two dimensions were nearly indistinguishable and that the Common Factor model may be a better representation of the covariance among externalizing phenotypes and SUDs. Nevertheless, we carried forward all three remaining models into the sensitivity and genetic correlation analyses to further evaluate this question.

### Sensitivity Analyses

#### Posthoc Models with Residual Correlations

Given that none of our models reached our thresholds of good fit, we ran the Common Factor, EXT-ARF, and BD-SUB models and included data-driven residual correlations among indicators. The primary purpose of these analyses was to determine the extent to which the parameter estimates in our primary models were influenced by inadequate fit. Briefly, factor loadings and correlations were stable across models with and without residual correlations (Supplementary Figure 1, Supplementary Table 5). The factor correlations between the externalizing and addiction risk factors (EXT-ARF model) were .90 and .88 in models with and without residual correlations, respectively. Factor correlations between the behavioral disinhibition and substance use factors (BD-SUB model) were .87 and .86 in models with and without residual correlations, respectively.

#### Variability of Factor Loadings

We first compared the loadings of each indicator across the Common Factor, EXT-ARF, and BD-SUB models. The loadings were relatively stable (Figure 3a), indicating that each indicator had a similar loading on its respective factor in each model. We next ran a series of sensitivity analyses to test the sensitivity of factors to their indicators by dropping one indicator from the model at a time. Figures 3b and 3c show the range of factor loadings and standard errors of an individual indicator when each other indicator was dropped from the model one at a time. On the whole, factors were not very sensitive to their indicators, as evidenced by the low variability of loadings (Figure 3b) and their standard errors (Figure 3c). We observed the most variability for the standard errors of SUDs in the EXT-ARF model. Factor correlations in the EXT-ARF and BD-SUB models were also stable, ranging from .78-.94 and .84-.90, respectively.

**Figure 3:**
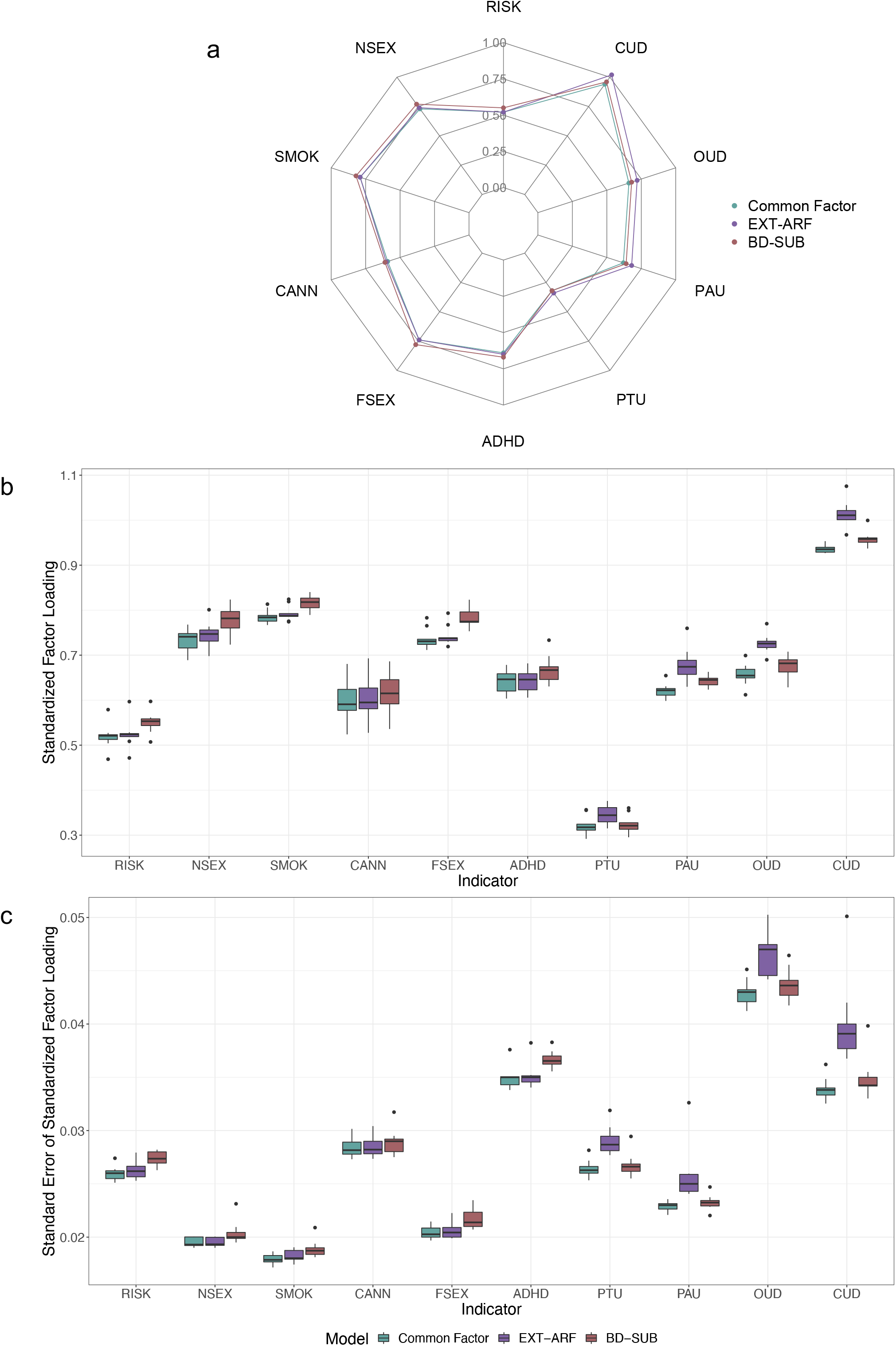
Variability of factor loadings across models. (a) Indicators’ loadings in the Common Factor, EXT-ARF, and BD-SUB Models. Variability of (b) factor loadings and (c) standard errors of the factor loadings of indicators when one indicator was dropped from the model at a time.

### Genetic Correlations

We estimated the genetic correlations between the factors in the Common Factor, EXT-ARF, and BD-SUB models and 84 preregistered phenotypes (see Figures 4-5 and Supplementary Table 6). We report results from models without residual correlations here and results for models with residual correlations in Supplementary Table 7.

**Figure 4.**
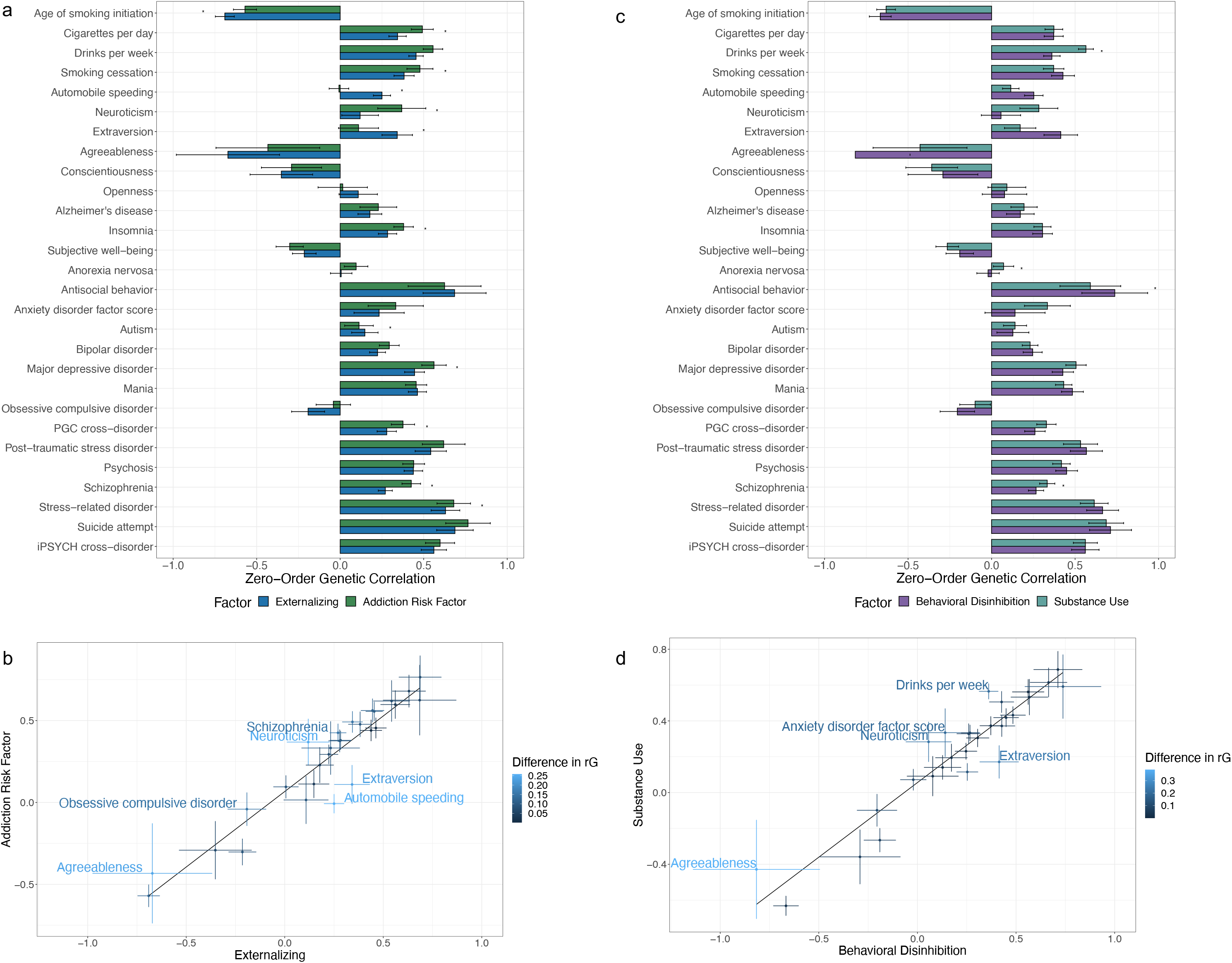
Comparison of genetic correlations of factors in the EXT-ARF and BD-SUB models with psychological, personality, and substance-use traits.

#### Are externalizing and SUDs differentially genetically correlated with other phenotypes?

The magnitude of the correlations of the EXT and ARF factors with external criteria were generally similar and their confidence intervals were often overlapping (Figure 4a). Further, the correlation between the effect sizes for the two factors was .96 and the median absolute difference in correlations was .06 (Figure 4b). The χ^2^ difference tests indicated that externalizing and the addiction risk factor were statistically significantly differentially correlated with 21 of 84 external criteria (see Figure 4a and Supplementary Table 6). However, many of these differences were small in magnitude, reflecting the sensitivity of the test to large sample size. The greatest differences EXT and ARF were observed for tobacco-related phenotypes (age of initiation [*r*_gARF_ = −.57, *r*_gEXT_ = −.69, |Δ*r*_g_| = .12] and cigarettes per day [*r*_gARF_ = .49, *r*_gEXT_ = .34, |Δ*r*_g_| = .15]); and certain personality traits (neuroticism [*r*_gARF_ = .37, *r*_gEXT_ = .12, |Δ*r*_g_|= .25] and extraversion [*r*_gARF_ = .11, *r*_gEXT_ = .34, |Δ*r*_g_| = .23]); and other forms of psychopathology (major depressive disorder [*r*_gARF_ = .56, *r*_gEXT_ = .45, |Δ*r*_g_| = .12 and schizophrenia [*r*_gARF_ = .43, *r*_gEXT_ = .27, |Δ*r*_g_| = .16).

#### Are behavioral disinhibition and substance use phenotypes differentially genetically correlated with other phenotypes?

A similar pattern emerged for the behavioral disinhibition and substance use factors in the BD-SUB model such that point estimates were similar and confidence intervals were often overlapping. χ^2^ difference tests indicated that 21 of 84 criteria were differentially associated with the behavioral disinhibition and substance use factors (see Figure 4c and Supplementary Table 6). The correlation between the effect sizes for the two factors was .95 (.01) and the median difference in correlations was .06 (Figure 4d). The greatest differences were observed for drinks per week (*r*_gBD_ = .36, *r*_gSUB_ = .57, |Δ*r*_g_| = .20), maternal smoking around birth (*r*_gBD_ = .65, *r*_gSUB_ = .83, |Δ*r*_g_| = .18), antisocial behavior (*r*_gBD_ = .74, *r*_gSUB_ = .59, |Δ*r*_g_| = .15).

#### With what is externalizing genetically correlated?

In Figure 5 and Supplemental Table 5, we report the correlations of the expanded externalizing factor that emerged from the Common Factor model with external criteria. Expanded externalizing was significantly genetically correlated with other psychiatric disorders, personality traits, socioeconomic, and substance use phenotypes. The strongest correlations were observed with drug exposure (*r*_g_ = .91 [.06]), prenatal tobacco exposure (*r*_g_ = .77 [.02]), the Townsend Index (*r*_g_ = .74 [.04]), suicide attempt (*r*_g_ = .73 [.05]), antisocial behavior (*r*_g_ = .69 [.07]), age of smoking initiation (*r*_g_ = −.68 [.03]), and agreeableness (*r*_g_ = −.62 [.14]).

**Figure 5.**
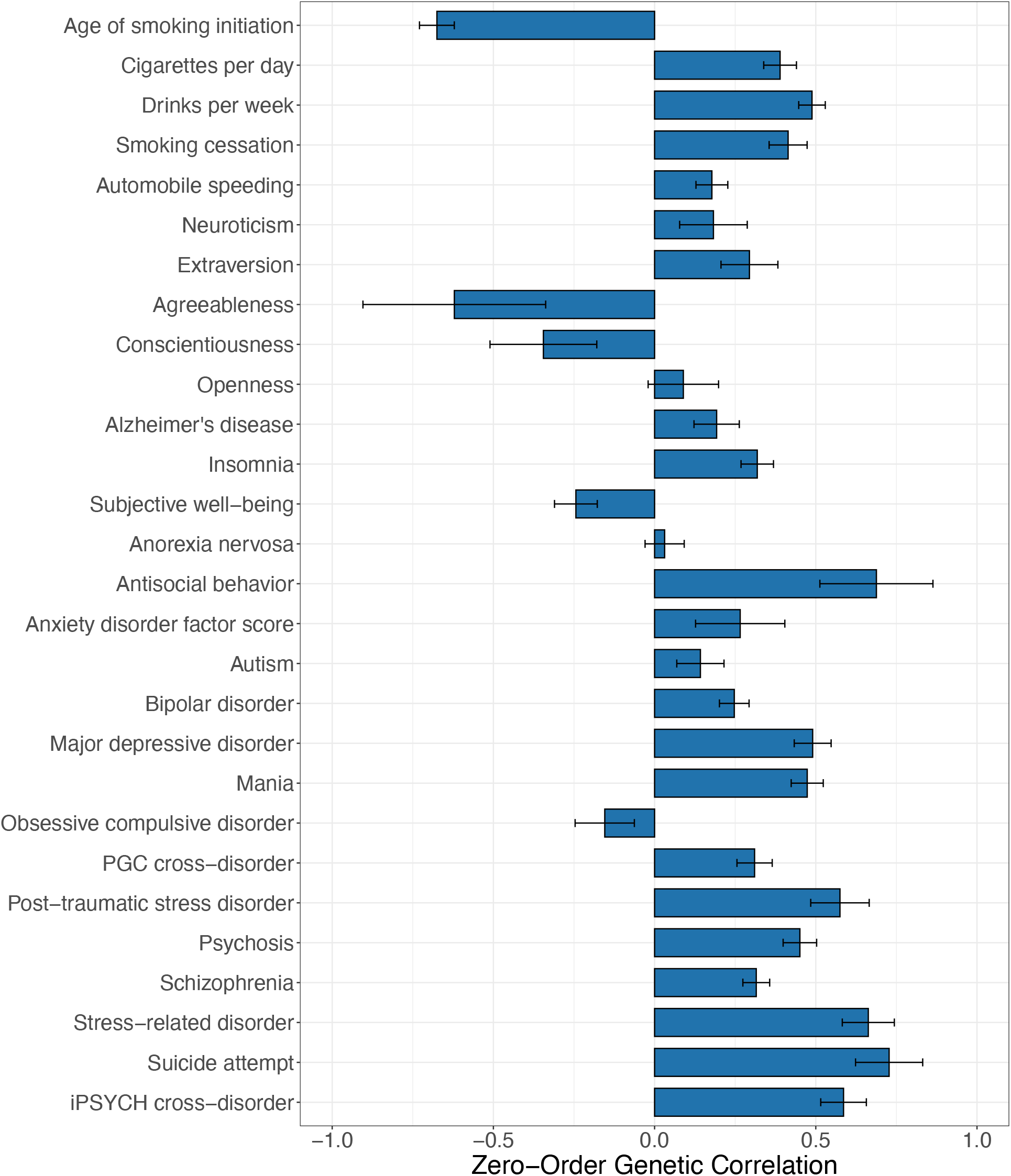
Genetic correlations between externalizing factor in the Common Factor model and psychological, personality, and substance-use traits.

#### Summary of Genetic Correlations

In each of the two-factor models (EXT-ARF and BD-SUB), the factors were similarly genetically correlated with external criteria such that the correlations of effect sizes were ≥ .95 and statistically significant differences were, for the most part, small in magnitude. This further supports the finding that, although model fit is improved in the two-factor models, the two dimensions do not represent meaningfully distinct dimensions of risk. The expanded externalizing factor in the Common Factor model was strongly genetically correlated with a variety of relevant psychological, substance use, and personality traits.

## Discussion

In this study, we investigated the structure of the shared genetic architecture among externalizing phenotypes and substance use disorders. To accomplish this, we tested a series of factor analytic models using Genomic SEM^22^. Although no model met all of our preregistered criteria for good git, our closest model assumed a common factor structure in which the genetic covariance among externalizing phenotypes and substance use disorders was explained by a single dimension of genetic risk. We settled on this model for two reasons: 1) the loadings in the common factor model were relatively homogeneous and the factor was well-represented by its indicators and 2) the high correlations between factors in the two correlated factors models as well as the similarity in the genetic correlations between these factors and external criteria suggest the factors are not meaningfully distinct. Further, the pattern of factor loadings and correlations was robust to the inclusion of residual correlations that improved model fit.

The finding that SUDs share a large proportion of variance with externalizing phenotypes is consistent with previous phenotypic SEM and twin literature. Many structural models of psychopathology include SUDs in the externalizing dimension^26,27^ and twin studies have found that genetic and environmental influences on SUDs largely operate through a broad externalizing factor^9,46^. Many structural models, however, place SUDs within specific subdimensions of externalizing, thus acknowledging variance that is unique to SUDs and not shared with all other forms of externalizing. For example, in the externalizing spectrum model^12^, substance use related traits are a subdimension of general externalizing and, in the HiTOP model^47^, SUDs are part of an externalizing subdimension representing disinhibited forms of externalizing, which are distinct from antagonistic forms. The number of well-powered GWAS of externalizing phenotypes may limit our ability to detect dimensionality in the current analyses and, as more GWAS of relevant traits become available, a more complex, hierarchical genetic architecture may be uncovered.

Results from this paper suggest that an expanded externalizing factor, which includes SUDs, can be used for future gene identification efforts. This is particularly important as gene identification efforts for SUDs have lagged behind consumption- and initiation-related traits and other forms of psychopathology, largely due to issues of power^48^. Simultaneous analysis of SUDs and genetically correlated traits, such as the externalizing phenotypes included in the current study, can boost power to detect associations for SUDs^22^. The use of Genomic SEM, in particular, also allows for the study of residual phenotypes (e.g., variance specific to cannabis use disorder unique from what it shares with other externalizing traits), which provides insights into genetic architecture that is specific to a given trait^49–51^. Pursuing these dual goals will allow us to overcome barriers to gene-identification in SUDs while still investigating any potential disorder-specific effects.

There were a few instances in which the externalizing and addiction risk factors were differentially associated with external criteria that warrant discussion. For example, age of smoking initiation was more strongly correlated with externalizing than with the addiction risk factor, which is consistent with evidence from twin studies suggesting that genetic variance for general externalizing, rather than variance specific to substance use, predicts age of onset^52^. The finding that the addiction risk factor is more strongly associated with neuroticism, major depression, and schizophrenia, and less strongly associated with extraversion, suggests that what distinguishes SUDs from externalizing is not necessarily something specific to risk for addiction, but rather risk shared with other forms of psychopathological distress.

This project is marked by two notable limitations. First, due to limited availability of GWAS summary statistics and technical issues involved with including multiple ancestries in a single Genomic SEM model, our analyses are limited to individuals of European ancestries. Second, our model does not account for other sources of genetic variance (e.g., internalizing and thought disorders) shared among externalizing and SUD phenotypes and which may contribute to the increased overlap among these phenotypes. These sources of variance may be especially important, given evidence that the addiction risk and externalizing factors are differentially associated with internalizing and thought disorder psychopathology.

In this study, we show that externalizing phenotypes and SUDs are influenced by a common dimension of genetic risk using Genomic SEM. We evaluated each of our models against a preregistered set of criteria and found that a common factor model best represented the relationships among our phenotypes. These results can be carried forward into future studies, in which we can leverage the genetic correlations among these phenotypes to boost power to detect associations for SUDs, the gene-identification efforts of which have lagged behind other psychiatric traits. This will also facilitate a more fine-grained exploration of the genetic influences unique to each externalizing phenotype, thereby allowing exploration of both broad and specific dimensions of genetic risk.

## Authors Contributions

HEP, PBB, and DMD were responsible for the original study concept and design. All authors provided revision of the preregistered analysis plan. HEP, PBB, and AH contributed to data acquisition and processing. HEP and TTM were responsible for data analysis and AH assisted with data analysis. HEP, AH, TTM, PBB, and DMD contributed to interpretation of findings. HEP drafted the original manuscript and all authors provided critical revision of the manuscript.

## Notes

https://osf.io/v8q2y/

### Competing Interest Statement

The authors have declared no competing interest.

## References

1 Florence, C. S., Zhou, C., Luo, F. & Xu, L. The Economic Burden of Prescription Opioid Overdose, Abuse, and Dependence in the United States, 2013. Medical Care 54 (2016).

2 Sacks, J. J., Gonzales, K. R., Bouchery, E. E., Tomedi, L. E. & Brewer, R. D. 2010 National and State Costs of Excessive Alcohol Consumption. American Journal of Preventive Medicine 49, e73–e79 (2015). https://doi.org/10.1016/j.amepre.2015.05.031

3 Louisa Degenhardt, F. C., Alize Ferrari, Damian Santomauro, Holly Erskine, Ana Mantilla-Herrara, Harvey Whiteford, Janni Leung, Mohsen Naghavi, Max Griswold, Jürgen Rehm, Wayne Hall, Benn Sartorius, James Scott, Stein Emil Vollset, Ann Kristin Knudsen, Josep Maria Haro, George Patton, Jacek Kopec, Deborah Carvalho Malta, Roman Topor-Madry, John McGrath, Juanita Haagsma, Peter Allebeck, Michael Phillips, Joshua Salomon, Simon Hay, Kyle Foreman, Stephen Lim, Ali Mokdad, Mari Smith, Emmanuela Gakidou, Christopher Murray, Theo Vos. The global burden of disease attributable to alcohol and drug use in 195 countries and territories, 1990-2016: a systematic analysis for the Global Burden of Disease Study 2016. The lancet. Psychiatry 5, 987–1012 (2018). https://doi.org:10.1016/S2215-0366(18)30337-7

4 Verhulst, B., Neale, M. C. & Kendler, K. S. The heritability of alcohol use disorders: a meta-analysis of twin and adoption studies. Psychol Med 45, 1061–1072 (2015). https://doi.org:10.1017/s0033291714002165

5 Verweij, K. J. et al. Genetic and environmental influences on cannabis use initiation and problematic use: a meta-analysis of twin studies. Addiction 105, 417–430 (2010). https://doi.org:10.1111/j.1360-0443.2009.02831.x

6 Kendler, K. S., Jacobson, K. C., Prescott, C. A. & Neale, M. C. Specificity of genetic and environmental risk factors for use and abuse/dependence of cannabis, cocaine, hallucinogens, sedatives, stimulants, and opiates in male twins. Am J Psychiatry 160, 687–695 (2003). https://doi.org:10.1176/appi.ajp.160.4.687

7 Kendler, K. S. & Myers, J. The boundaries of the internalizing and externalizing genetic spectra in men and women. Psychol Med 44, 647–655 (2014). https://doi.org:10.1017/S0033291713000585

8 Barr, P. B. & Dick, D. M. The Genetics of Externalizing Problems. Curr Top Behav Neurosci 47, 93–112 (2020). https://doi.org:10.1007/7854_2019_120

9 Krueger et al. Etiological Connections among Substance Dependence, Antisocial Behavior, and Personality: Modeling the Externalizing Spectrum. Journal of Abnormal Psychology 111, 411–424 (2002). https://doi.org:10.1037/0021-843X.111.3.411

10 Watts, A. L., Poore, H. E. & Waldman, I. D. Riskier Tests of the Validity of the Bifactor Model of Psychopathology. Clinical Psychological Science 7, 1285–1303 (2019). https://doi.org:10.1177/2167702619855035

11 Blanco, C. et al. The Space of Common Psychiatric Disorders in Adolescents: Comorbidity Structure and Individual Latent Liabilities. Journal of the American Academy of Child & Adolescent Psychiatry 54, 45–52 (2015). https://doi.org/10.1016/j.jaac.2014.10.007

12 Krueger, Markon, Patrick, Benning & Kramer. Linking Antisocial Behavior, Substance Use, and Personality: An Integrative Quantitative Model of the Adult Externalizing Spectrum. Journal of Abnormal Psychology 116, 645–666 (2007). https://doi.org:10.1037/0021-843X.116.4.645

13 Karlsson Linnér, R. et al. Multivariate analysis of 1.5 million people identifies genetic associations with traits related to self-regulation and addiction. Nature Neuroscience 24, 1367–1376 (2021). https://doi.org:10.1038/s41593-021-00908-3

14 Card, N. A. & Little, T. D. Proactive and reactive aggression in childhood and adolescence: A meta-analysis of differential relations with psychosocial adjustment. International Journal of Behavioral Development 30, 466–480 (2006). https://doi.org:10.1177/0165025406071904

15 Tackett, J. L., Daoud, S. L., De Bolle, M. & Burt, S. A. Is relational aggression part of the externalizing spectrum? a bifactor model of youth antisocial behavior. Aggress Behav 39, 149–159 (2013). https://doi.org:10.1002/ab.21466

16 Kotov, R., Gamez, W., Schmidt, F. & Watson, D. Linking “big” personality traits to anxiety, depressive, and substance use disorders: a meta-analysis. Psychol Bull 136, 768–821 (2010). https://doi.org:10.1037/a0020327

17 Watts, A. L., Poore, H. E., Lilienfeld, S. O. & Waldman, I. D. Clarifying the associations between Big Five personality domains and higher-order psychopathology dimensions in youth. Journal of Research in Personality 82 (2019). https://doi.org:10.1016/j.jrp.2019.07.002

18 Walters, R. K. et al. Transancestral GWAS of alcohol dependence reveals common genetic underpinnings with psychiatric disorders. Nat Neurosci 21, 1656–1669 (2018). https://doi.org:10.1038/s41593-018-0275-1

19 Waldman, I. D., Poore, H. E., Luningham, J. M. & Yang, J. Testing structural models of psychopathology at the genomic level. World Psychiatry 19, 350–359 (2020). https://doi.org:10.1002/wps.20772

20 Sanchez-Roige, S. et al. Genome-Wide Association Study Meta-Analysis of the Alcohol Use Disorders Identification Test (AUDIT) in Two Population-Based Cohorts. Am J Psychiatry 176, 107–118 (2019). https://doi.org:10.1176/appi.ajp.2018.18040369

21 Johnson, E. C. et al. A large-scale genome-wide association study meta-analysis of cannabis use disorder. Lancet Psychiatry 7, 1032–1045 (2020). https://doi.org:10.1016/s2215-0366(20)30339-4

22 Grotzinger, A. D. et al. Genomic structural equation modelling provides insights into the multivariate genetic architecture of complex traits. Nat Hum Behav 3, 513–525 (2019). https://doi.org:10.1038/s41562-019-0566-x

23 Bulik-Sullivan, B. K. et al. LD Score regression distinguishes confounding from polygenicity in genome-wide association studies. Nat Genet 47, 291–295 (2015). https://doi.org:10.1038/ng.3211

24 Hatoum, A. S. et al. The addiction risk factor: A unitary genetic vulnerability characterizes substance use disorders and their associations with common correlates. Neuropsychopharmacology (2021). https://doi.org:10.1038/s41386-021-01209-w

25 Hatoum, A. S. et al. Multivariate genome-wide association meta-analysis of over 1 million subjects identifies loci underlying multiple substance use disorders. medRxiv, 2022.2001.2006.22268753 (2022). https://doi.org:10.1101/2022.01.06.22268753

26 Forbes, M. K. et al. Three recommendations based on a comparison of the reliability and validity of the predominant models used in research on the empirical structure of psychopathology. J Abnorm Psychol 130, 297–317 (2021). https://doi.org:10.1037/abn0000533

27 Lahey, B. B., Krueger, R. F., Rathouz, P. J., Waldman, I. D. & Zald, D. H. A hierarchical causal taxonomy of psychopathology across the life span. Psychol Bull 143, 142–186 (2017). https://doi.org:10.1037/bul0000069

28 Greene, A. L. & Eaton, N. R. The temporal stability of the bifactor model of comorbidity: An examination of moderated continuity pathways. Compr Psychiatry 72, 74–82 (2017). https://doi.org:10.1016/j.comppsych.2016.09.010

29 Keyes, K. M. et al. Thought disorder in the meta-structure of psychopathology. Psychological Medicine 43, 1673–1683 (2013). https://doi.org:10.1017/S0033291712002292

30 Castellanos-Ryan, N. et al. The structure of psychopathology in adolescence and its common personality and cognitive correlates. J Abnorm Psychol 125, 1039–1052 (2016). https://doi.org:10.1037/abn0000193

31 Verona, E., Javdani, S. & Sprague, J. Comparing factor structures of adolescent psychopathology. Psychol Assess 23, 545–551 (2011). https://doi.org:10.1037/a0022055

32 Vollebergh, W. A. et al. The structure and stability of common mental disorders: the NEMESIS study. Arch Gen Psychiatry 58, 597–603 (2001). https://doi.org:10.1001/archpsyc.58.6.597

33 Karlsson Linnér, R. et al. Genome-wide association analyses of risk tolerance and risky behaviors in over 1 million individuals identify hundreds of loci and shared genetic influences. Nature Genetics 51, 245–257 (2019). https://doi.org:10.1038/s41588-018-0309-3

34 Demontis, D. et al. Discovery of the first genome-wide significant risk loci for attention deficit/hyperactivity disorder. Nat Genet 51, 63–75 (2019). https://doi.org:10.1038/s41588-018-0269-7

35 Liu, M. et al. Association studies of up to 1.2 million individuals yield new insights into the genetic etiology of tobacco and alcohol use. Nature Genetics 51, 237–244 (2019). https://doi.org:10.1038/s41588-018-0307-5

36 Pasman, J. A. et al. GWAS of lifetime cannabis use reveals new risk loci, genetic overlap with psychiatric traits, and a causal influence of schizophrenia. Nat Neurosci 21, 1161–1170 (2018). https://doi.org:10.1038/s41593-018-0206-1

37 Zhou, H. et al. Genome-wide meta-analysis of problematic alcohol use in 435,563 individuals yields insights into biology and relationships with other traits. Nat Neurosci 23, 809–818 (2020). https://doi.org:10.1038/s41593-020-0643-5

38 Zhou, H. et al. Association of OPRM1 Functional Coding Variant With Opioid Use Disorder: A Genome-Wide Association Study. JAMA Psychiatry 77, 1072–1080 (2020). https://doi.org:10.1001/jamapsychiatry.2020.1206

39 Schoeler, T. et al. Novel biological insights into the common heritable liability to substance involvement: a multivariate genome-wide association study. Biological Psychiatry (2022). https://doi.org/10.1016/j.biopsych.2022.07.027

40 Hu, L. t. & Bentler, P. M. Cutoff criteria for fit indexes in covariance structure analysis: Conventional criteria versus new alternatives. Structural Equation Modeling: A Multidisciplinary Journal 6, 1–55 (1999). https://doi.org:10.1080/10705519909540118

41 Waldman, I. D. et al. Recommendations for Adjudicating Among Alternative Structural Models of Psychopathology. Clinical Psychological Science (*in press*).

42 Graham, J. M. Congeneric and (Essentially) Tau-Equivalent Estimates of Score Reliability: What They Are and How to Use Them. Educational and Psychological Measurement 66, 930–944 (2006). https://doi.org:10.1177/0013164406288165

43 Hancock, G. R. & Mueller, R. O. in Structural equation modeling: Present and future—A Festschrift in honor of Karl Jöreskog (eds R. Cudeck, S. du Toit, & D. Sorbom) 195–216 (Scientific Software International, 2001).

44 Revelle, W. & Zinbarg, R. E. Coefficients Alpha, Beta, Omega, and the glb: Comments on Sijtsma. Psychometrika 74, 145 (2009). https://doi.org:10.1007/s11336-008-9102-z

45 Team, R. C. R: A language and environment for statistical computing. (R Foundation for Statistical Computing, 2022).

46 Young, Stallings, Corley, Krauter & Hewitt. Genetic and Environmental Influences on Behavioral Disinhibition. American Jounral of Medical Genetic 96, 684–695 (2000). https://doi.org:10.1002/1096-8628

47 Krueger, R. F. et al. Validity and utility of Hierarchical Taxonomy of Psychopathology (HiTOP): II. Externalizing superspectrum. World Psychiatry 20, 171–193 (2021). https://doi.org/10.1002/wps.20844

48 Deak, J. D. & Johnson, E. C. Genetics of substance use disorders: a review. Psychological Medicine 51, 2189–2200 (2021). https://doi.org:10.1017/S0033291721000969

49 Demange, P. A. et al. Investigating the genetic architecture of noncognitive skills using GWAS-by-subtraction. Nature Genetics 53, 35–44 (2021). https://doi.org:10.1038/s41588-020-00754-2

50 Barr, P. B. et al. Parsing Genetically Influenced Risk Pathways: Genetic Loci Impact Problematic Alcohol Use Via Externalizing and Specific Risk. medRxiv, 2021.2007.2020.21260861 (2021). https://doi.org:10.1101/2021.07.20.21260861

51 Mallard, T. T. et al. Item-Level Genome-Wide Association Study of the Alcohol Use Disorders Identification Test in Three Population-Based Cohorts. Am J Psychiatry 179, 58–70 (2022). https://doi.org:10.1176/appi.ajp.2020.20091390

52 Dick, D. M., Adkins, A. E. & Kuo, S. I. C. Genetic influences on adolescent behavior. Neuroscience & Biobehavioral Reviews 70, 198–205 (2016). https://doi.org/10.1016/j.neubiorev.2016.07.007

