## Supplementary Figure 1 for "A Multivariate Approach to Understanding the Genetic Overlap between Externalizing Phenotypes and Substance Use Disorders"

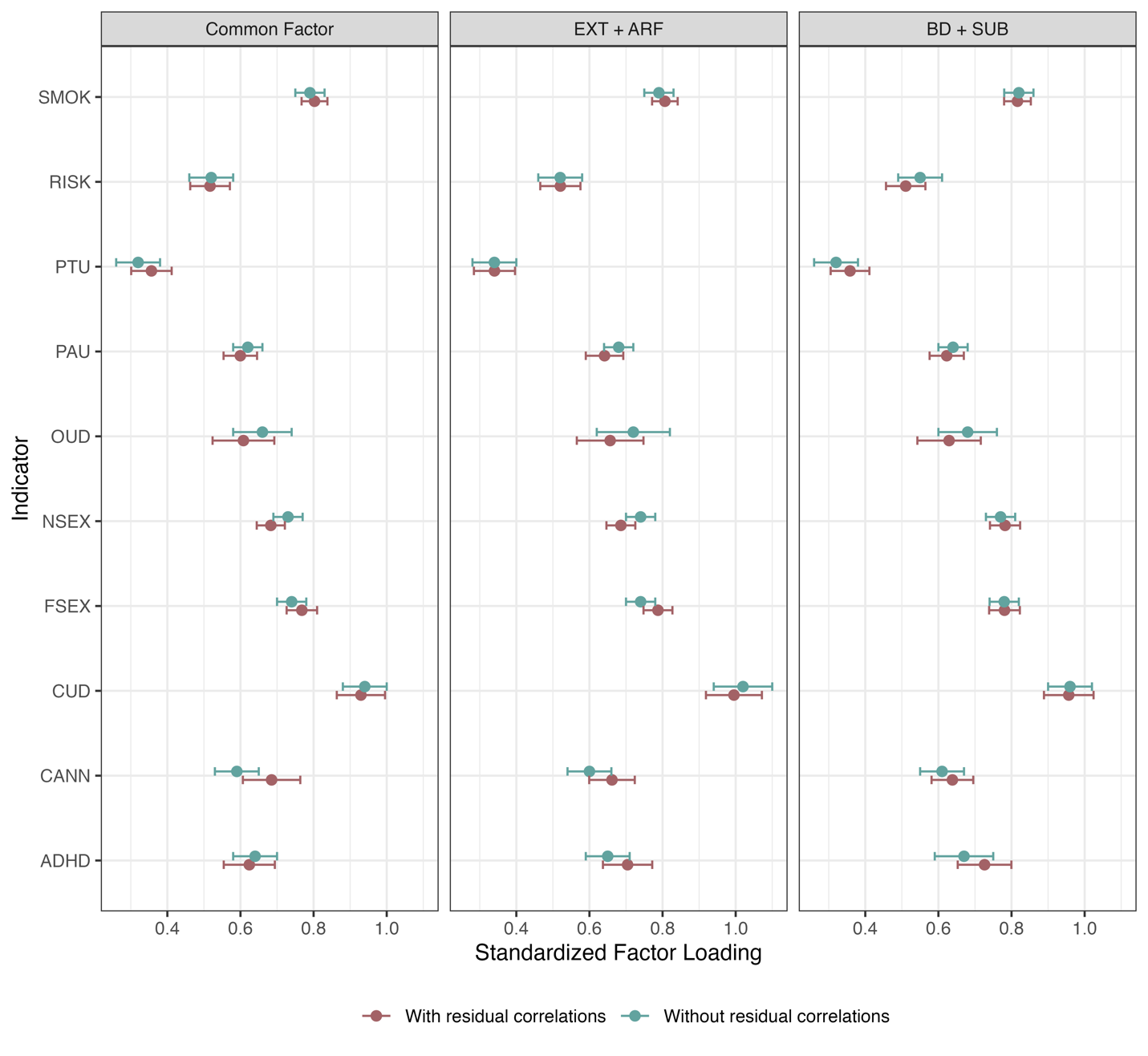

Supplementary Figure 1. Indicators’ factor loadings in models with and without data-driven factor loadings.
